# Transcranial electric stimulation modulates firing rate at clinically relevant intensities

**DOI:** 10.1101/2023.11.24.568618

**Authors:** Forouzan Farahani, Niranjan Khadka, Lucas C. Parra, Marom Bikson, Mihály Vöröslakos

## Abstract

Notwithstanding advances with low-intensity transcranial electrical stimulation (TES), there remain questions about the efficacy of clinically realistic electric fields on neuronal function. We used Neuropixels 2.0 probe with 384 channels in an in-vivo rat model of TES to detect effects of weak fields on neuronal firing rate. High-density field mapping and computational models verified field intensity (1 V/m in hippocampus per 50 µA of applied skull currents). We demonstrate that electric fields below 0.5 V/m acutely modulate firing rate in 5% of neurons recorded in the hippocampus. At these intensities, average firing rate effects increased monotonically with electric field intensity at a rate of 7 % per V/m. For the majority of excitatory neurons, firing increased for cathodal stimulation and diminished for anodal stimulation. While more diverse, the response of inhibitory neurons followed a similar pattern on average, likely as a result of excitatory drive. Our results indicate that responses to TES at clinically relevant intensities are driven by a fraction of high-responder excitatory neurons, with polarity-specific effects. We conclude that transcranial electric stimulation is an effective neuromodulator at clinically realistic intensities.

## Introduction

The effects of transcranial electric stimulation on neural activity in the brain have been known since the 1960^1–3^. The acute effects on neuronal firing rate are particularly well established. Namely, the electric fields generated within the brain by transcranial current stimulation can incrementally polarize cell membranes^4^ and thus modulate ongoing cell firing^5,6^. The effect acts at the time scale of the neuronal membrane (∼30ms) and thus is relevant for direct current (DC) and most effective for alternating currents (AC) of 30 Hz or less^7,8^. This acute neuromodulatory effect can be predicted from the orientation and intensity of local electric fields^9^. These cellular mechanisms established with in vitro animal experiments, also point to network effects^10,11^, which can be properly studied only in the intact brain.

However, despite numerous in-vivo animal studies in the intervening decades^12–24^, there is still a lack of clarity as to whether the effects observed are clinically relevant, for one simple reason: in vivo animal experiments have not adequately characterized electric field magnitudes in the brain. In particular, a significant gap has emerged^25^ between electric fields measured in vivo in the human brain, which are at or below 0.5 V/m^15,26,27^ and field intensities used for in vitro animal experiments, which are mostly at or above 5 V/m^28^. Thus, it is difficult to interpret and link results from in vivo animal experiments to cellular effects observed in vitro. Nor is it clear that the in vivo animal experiments have any relevance to the behavioral effects observed in human clinical studies.

To close this gap, we measured here for the first-time electric fields magnitude and their effects on neuronal firing rate in vivo in rats and established calibrated computational models of current flow. To do so, we first calibrated our recording equipment on a phantom, and performed in vivo field measurements in cortex and hippocampus in a rodent TES model. Then, using high-channel probes (Neuropixels2.0)^29^ we analyzed firing rate of individual putative pyramidal and interneurons in response to short (2s) DC stimulation. We demonstrate here acute modulation of neuronal firing rate with 0.5 V/m electric fields. Polarity-specific sensitivity at such low fields were governed by a small population of excitatory neurons. Prior studies have shown that changes in a small number of neurons can lead to behavioral effects^30,31^. Thus, clinically relevant TES intensities produce neuronal firing changes sufficient, in principle, to impact human brain function.

## Results

### Measurement and modeling of TES-induced electric fields in motor cortex of rats

To characterize the effects of TES it is necessary to properly calibrate electric field measurements, which is the main determining factor for acute effects on neuronal function^32^. After characterizing our stimulation and recording system using agar phantom (**Suppl. Fig. 1**), we measured field intensity intracranially and built an anatomically detailed computational model of our electrode montage. In our experimental setup we applied sinusoidal alternating current (10,100 and 1000 Hz) in two anesthetized rats. To electrically isolate the animal from the metallic stereotactic frame, we 3D-printed a non-conducting nose holder and ear bars (**Fig. 1a and Suppl. Fig. 2**, Clear V4 resin, Formlabs) and placed the animal on a non-conducting surface. A platinum electrode was affixed to the skull over the forelimb motor cortex (1.5 mm anterior to bregma and 3 mm lateral from midline) within a chamber loaded with conductive gel. The pocket to hold gel and TES electrode was made of dental cement (**Fig. 1b**, 3 by 3 mm, GC Unifast). The return electrode was a platinum mesh (10 by 10 mm) implanted in the chest wall^19^ (**Fig. 1a**). This electrode montage provided electro-chemical stability and free range of movements in behaving rats. To measure the electric field generated by transcranial stimulation, we used a multi-channel, custom-built recording electrode matrix (n = 4 channels in total, 2 channels per shank, 1 mm distance between shanks and channels, **Fig. 1b and Suppl. Fig. 3**). After a craniotomy through the parietal bone, we inserted the electrode matrix into the motor cortex from the lateral side and sealed it with non-conductive silicon (Suppl. Fig. 2b, Kwik Cast silicone, Kwik-Cast). We found that electric field magnitude increased linearly with stimulation current, with similar slope at the three stimulation frequencies (Fig. 2c, slope: 15.0 V/m/µA). In a second animal we measured fields of twice this magnitude (not shown, slope 30.0 V/m).

**Figure 1.**
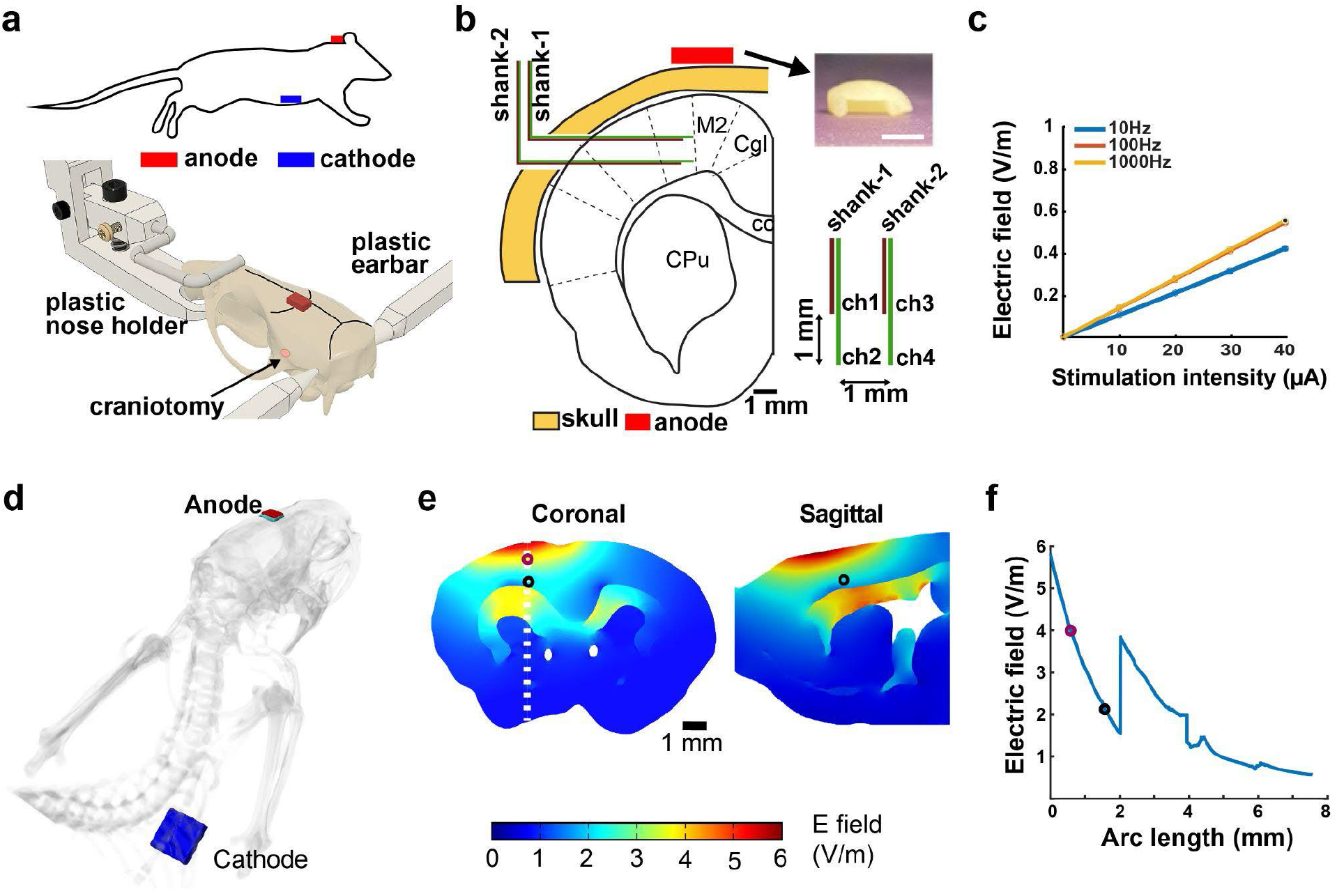
Measurement and modeling of TES-induced electric field in motor cortex. **a)** Electric field measurement in the motor cortex of rats. Top: anode is affixed to the skull above the primary motor cortex (3 by 3 mm platinum plate), cathode is implanted inside the chest wall (10 by 10 mm platinum mesh). Bottom: 3D-printed nose holders and ear bars are used to isolate the animal from the metallic components of the stereotactic frame during measurements. The rat skull is shown inside the nose holder with an attached anode (red rectangle) and craniotomy in the parietal bone. **b)** Schematic of the position of recording electrodes in the motor cortex. Note that electrodes were inserted from the lateral side of the skull through the temporal craniotomy. Top, right: Customized holder for the stimulation electrode (scale bar is 3 mm). Bottom, right: schematic of the custom-built, 2-shank, 4-channel tungsten recording matrix. Each shank had 2 recording channels (both ch1 and ch2 and shank-1 and shank-2 are separated by 1 mm). **c)** Increasing stimulation intensity induces an increasing electric field in the motor cortex . **d)** Anatomically accurate FEM model including anode (red) and gel (green) placed on the skull and cathode implanted in the chest (blue). **e)** Distribution of field magnitude estimated with the current flow model at 150 μA current. **f)** Field amplitude as a function of distance from the cortical surface moving in radial direction (Arc length). The discontinuity is due to a discontinuity in conductivity (white matter of corpus colosseum has lower conductivity than gray matter, 0.126 S/m vs 0.276 S/m)

**Figure 2.**
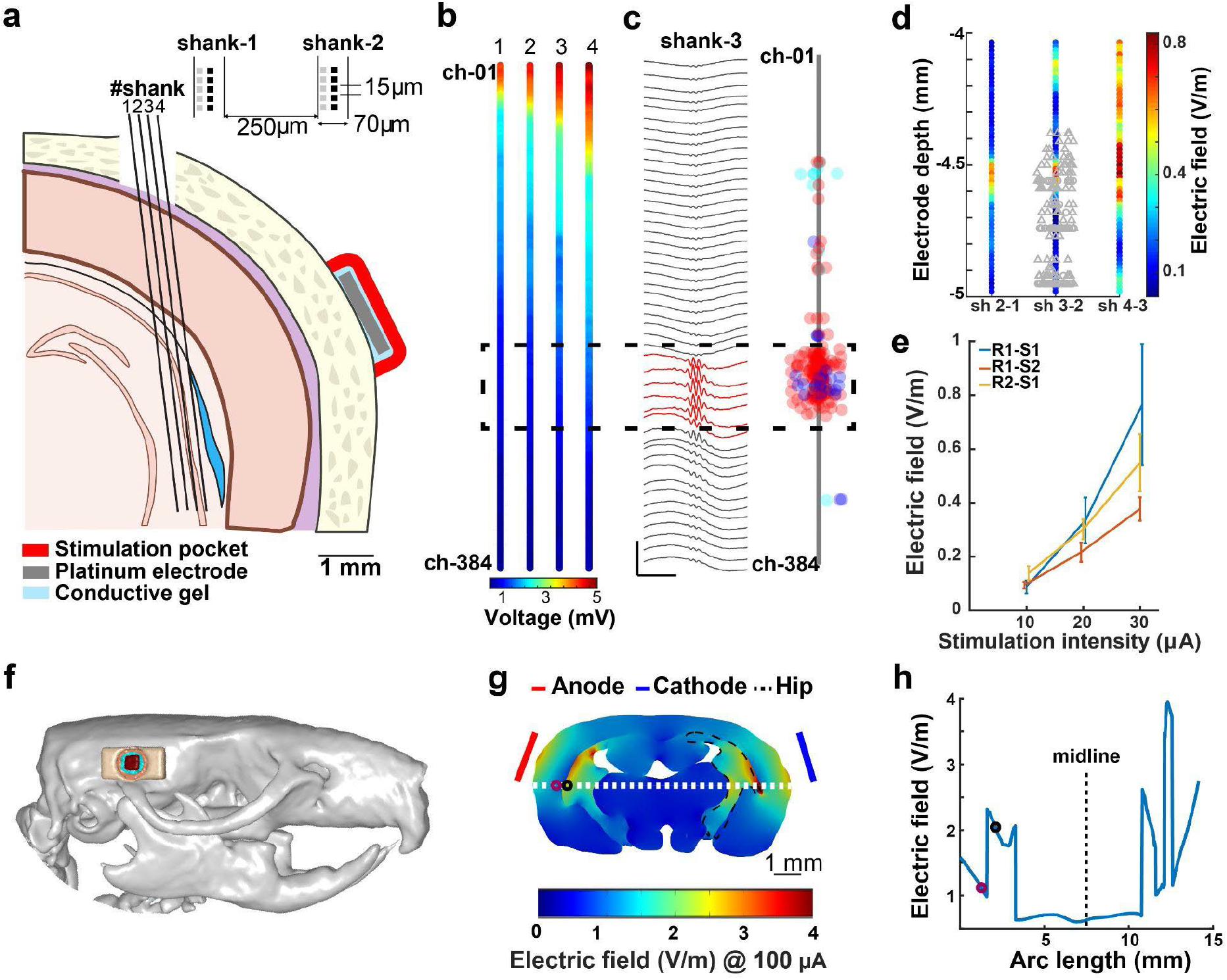
Measurement and modeling of TES-induced electric field in hippocampus. **a)** Electric field measurement in the hippocampus of freely moving rats. Anode and cathode are placed on the temporal bone (2 by 2 mm platinum plate). Multi-shank, multi-site silicon probe is used to measure the electric field (probe is inserted at 10 degrees), details of the shanks are shown on the right. **b)** TES-induced (30 µA, 100 Hz) peak-to-peak voltage changes measured in 4 shanks. Colors indicate the peak-to-peak average measured on each channel (n = 500 repetitions, n = 384 channels/shank, recorded sequentially from n = 4 shanks). Note the increasing voltage values closer to the stimulation electrode (shank-4). **c)** Localizing cellular layer of hippocampus using electrophysiological markers. Left: ripple triggered average LFP traces recorded on shank-3 linear configuration (n = 48 channels, every 8^th^ channel is shown). Red channels show the location of the maximum ripple amplitude. Right: schematic of shank-3 is shown with the putative location of recorded neuron somata (n = 181 putative pyramidal cells, 81 narrow interneurons and 2 wide interneurons, red, blue, and cyan circles, respectively). Single units were clustered in the cellular layers of the hippocampus (ch-01 represents brain surface). **d)** TES-induced electric fields recorded in the cellular layer of the hippocampus (black dashed rectangle, n = 64 channels per shank). The location of recorded neuron somas is overlaid in gray on shank 3-2. **e)** Increasing stimulation intensity (10, 20 and 30 µA) induces increasing intracerebral electric field (0.1, 0.28 and 0.56 V/m, R = 0.75, p < 0.001). **f)** Electrode montage in the rat model. **g)** Modeling results of TES-induced electric fields in the coronal plane (4.8 mm posterior from bregma) at 100µA. **h)** Electric field intensity along the white dotted line in panel g. The discontinuity in electric field is due to discontinuity in conductance between white and gray matter (see **Suppl. Fig. 4**)

**Figure 3.**
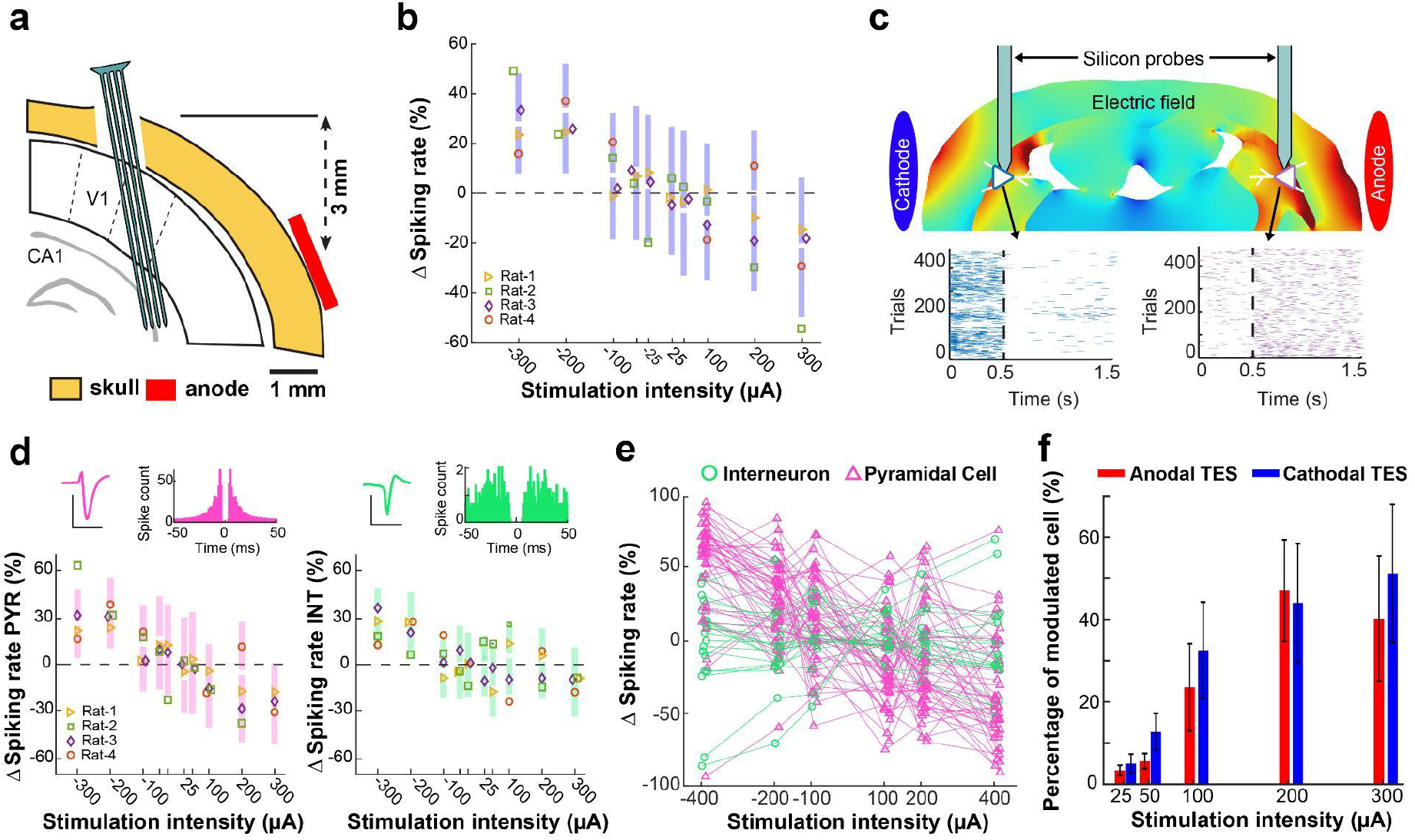
Electric field dependent change of firing rate of hippocampal neurons. **a)** Schematic of experimental setup. Multi-shank, multi-site silicon probe is used to measure neuronal activity in the intermediate CA2. **b)** TES induced a polarity and intensity dependent modulation of neuronal firing in the hippocampus (R = −0.33, P < 0.001, n = 510 neurons in 4 rats). **c)** Response of two putative pyramidal cells recorded from both hippocampi simultaneously using two 32-channel silicon probes. Blue and purple triangle shows the location of the cells’ somata overlaid on the electric field model. The neuron closer to the cathode (blue neuron) was excited by the stimulation as shown by the peristimulus time histogram. The neuron closer to the anode showed an opposite response (purple neuron). **d)** Recorded neurons are classified into putative excitatory (top, left) and putative inhibitory neurons (top, right) based on their waveform and autocorrelation histogram. The scale bar is 0.1 mV and 1 ms. Bottom: TES influenced the spiking rate of both putative pyramidal cells (bottom, left, R = −0.34, p < 0.001, n = 359 neurons in 4 rats) and putative interneurons (bottom, right, R = −0.3, p < 0.001, n = 151 neurons in 4 rats). **e)** Some neurons were modulated in the opposite direction as the average response of the hippocampus. Note the three cells that were inhibited by cathodal and excited by anodal TES (example session from one rat). **f)** Change in percentage of significantly modulated neurons (Wilcoxon signed rank test p<0.05) by cathodal (blue) and anodal (red) stimulation (change in spiking activity is measured relative to the 3-second stimulation free period between 3-second stimulation epochs). Note, the number of modulated cells increased with intensity but were similar across TES polarities (bar graphs show the mean, error bars represent the SEM, n = 4 rats).

To build the computation model, we used a high-resolution (0.1 mm) MRI (magnetic resonance image) of a healthy rat, which had been segmented into tissue masks and assigned each a conductivity value (skull – 0.02 S/m, cerebrospinal fluid – 1.7 S/m, gray matter – 0.276 S/m, white matter – 0.126 S/m, hippocampus – 0.126 S/m) based on prior work^37,38^. We generated a Volumetric Finite Element method (FEM) model using Simpleware (Synopsys Inc., CA). The resulting volumetric meshes were later imported into COMSOL Multiphysics 5.5 (COMSOL Inc., MA, USA) to generate FEM models and solved for electric fields under steady-state assumption. We simulated our experimental setup by placing electrodes in the model on the skull above the motor cortex (anode, 3×3mm) and intercostal muscles (cathode, 8 by 8 mm) (**Fig 1d**). We applied constant current density through one electrode (anode: 150 µA) while grounding the other electrode (cathode). The external boundaries were electrically insulated (J.n = 0). Corresponding voltage and electric fields were quantified from the simulations. At the motor cortex location corresponding to the in vivo field recordings (Fig. 2e, circle) the model estimates an electric field of 2.26 V/m (**Fig. 1e**). This corresponds to 15.07 V/m per mA and is within the range measured in-vivo. Although it should be noted that there is a strong gradient as one moves radially (**Fig. 1f**) - moving just 1mm closer to the stimulating electrode the electric field per applied current doubles to 30 V/m per mA - and the recording matrix has 1 mm side length. The model indicated that only one hemisphere was affected by TES using our electrode montage (**Fig. 1e**).

### Measurement and modeling of TES-induced electric fields in hippocampus of rats

The field measurements and model established that 100 µA stimulation can induce 1.5-3 V/m fields in motor areas. The exact field magnitude strongly depends on the recording location and thus, it has to be measured in the precise region of interest. We were interested in neural responses in the hippocampus, and so we decided to measure fields again with the same electrodes we will use for neural activity. We implanted Neuropixels (NP) 2.0 probes^29^ in the intermediate CA2 region of freely moving rats (**Fig. 2a**, n = 2 rats, 4.8 mm posterior to bregma and 4.6 mm lateral to midline, angled at 10 degrees). We applied electrical current through two skull electrodes (2 mm by 2mm platinum plates), but this time affixed to the temporal bone bilaterally (**Fig. 2a**). We took advantage of the 5120 contacts available on the NP 2.0 probe, to select 384 channels for recording from each shank. We chose a single shank, linear configuration spanning 5760 µm (15 µm separation per channel) to record electric potential during sinusoidal TES (100 Hz, n = 500 cycles, at 10, 20, and 30 µA intensity) sequentially from each of the 4 implanted shanks. (**Fig. 2b**). As expected, we recorded higher voltages on the most lateral shank (**Fig. 2b**, shank-4 was closest to the stimulation electrode, each shank is separated by 250 µm). Additionally, we recorded higher voltage values following the curvature of the brain surface which likely reflects shunting caused by cerebrospinal fluid in the meningeal space^27,39,40^. To measure the electric fields in the hippocampus, we first localized the cellular layer of CA2 using electrophysiological markers (**Fig. 2c**). We detected sharp wave ripples in the local field potential (LFP) signal and calculated the ripple triggered average signal across 48 channels (**Fig. 2c, left**; 12-channel steps corresponding to 180 µm inter-site distance) and we identified the channel with maximum ripple power (**Fig 2c, left**; highlighted channels in red). We determined the position of individual neuronal somata using spike sorting and spike-amplitude trilateration (**Fig. 2c, right**; ^41,42^). To calculate the hippocampal electric field, we used ± 32 channels around the center of these soma locations (**Fig. 2d**). Similar to the motor cortex, we found that increasing stimulation intensity (10, 20 and 30 µA) induced increasing intracerebral electric fields (0.1 ± 0.01, 0.28 ± 0.03 and 0.56 ± 0.11 V/m, mean ± SEM, n = 3 sessions from 2 rats, R = 0.76, p < 0.001). This corresponds to 10, 14 and 18.7 V/m per mA and thus somewhat less than the cortical measures, as expected. To simulate our experimental setup, we placed electrodes over the parietal bone (**Fig 2f**). We applied 100 µA current through one electrode (anode) while grounding the other electrode (cathode). The model predicted an electric field of 2.1 V/m in white matter and 1.2 V/m in gray matter in the hippocampus (**Fig. 2g, h**). This corresponds to 12-21 V/m per mA of applied current and is consistent with what we observed in the experimental recording above. As expected, the magnitude of the electric field dropped with distance from the cortex but increased at the boundary of white-gray matter transition (**Fig. 2g, h**).

### Intensity and polarity dependent effects of single unit activity induced by TES

Single-unit action potentials – which are not always available in animals and are rarely possible in humans – are the most direct measurement of neural activity. We quantified how different TES intensities (25 to 400 µA) can affect the spiking activity of neurons in the hippocampus. These currents generate fields in the range of 0.375 V/m to 6 V/m assuming the observed 15 V/m per mA applied. We performed these measurements using the same rats where we measured electric fields and using the same recording and stimulation electrodes. TES was applied for 4 seconds and repeated 400 times with 4 seconds intervals of no stimulation. Single-unit activity was recorded from the CA2 region (**Fig. 3a, Suppl. Fig. 5**) in 4 rats freely moving in their home cage, and one anesthetized rat (**Suppl. Table 1**)^29,43^. Depending on the polarity of the stimulation, putative single units either increased (cathodal TES) or decreased (anodal TES) their spiking activity (**Fig. 3b**; slope: −3.75% per V/m, R = −0.33, P < 0.001, n = 510 neurons). Mean percent change in firing rate (FR) of neurons was 24.57 ± 1.53 (−300 µA), 27.65 ± 1.51 (−200 µA), 4.83 ± 1.61 (−100 µA), 7.3 ± 2.03 (−50 µA), 6.18 ± 1.87 (−25 µA), −2.5 ± 1.92 (25 µA), −3 ± 2.08 (50 µA), −6.64 ± 1.74 (100 µA), −7.69 ± 1.94 (200 µA), and −19.82 ±1.82 (300 µA; **Fig. 3b**, mean ± SEM, n = 510 neurons in 4 rats and n = 394 neurons in 3 rats). We have tested higher intensities in a urethane anesthetized rat and found that the effects did not saturate. Specifically, cathodal stimulation increased the spiking rate by 37.22 ± 5.13% (−400 µA), while anodal TES further decreased the activity of neurons by −25.4 ± 4.51% (400 µA, n = 68 neurons, **Suppl. Fig. 6**). To confirm the opposing effect on spiking activity of hippocampal cells underneath the anode and cathode, we recorded from both hippocampi simultaneously using two, 32-channel silicon probes in an anesthetized rat^15^. Our modeling results anticipated that the electric field’s magnitude would be comparable in both hemispheres, but with opposing orientation relative to the orientation of pyramidal neurons. Neurons under the cathode were excited (**Fig. 3c**, blue neuron), whereas those under the anode were inhibited during TES (**Fig. 3c**, purple neuron, n = 400 trials, 500 ms stimulation followed by 1 s stimulation free epochs). This is the expected direction of effects given that hippocampal pyramidal neurons have the opposite orientation to cortical-surface neurons and therefore radially outward currents are soma-depolarizing for hippocampal neurons^4,44^.

In the freely behaving animals, single units were classified into putative pyramidal cell and interneuron types based on waveform and spike train characteristics (**Figure 3d**, top; see Methods). Stimulation exerted clear and predictable effects on the spiking rate of putative pyramidal cells (**Fig. 3d**, left; R = −0.34, p < 0.001, n = 359 putative pyramidal cells) and putative interneurons (**Fig. 3d**, right; R = −0.3, p < 0.001, n = 151 putative interneurons). A linear fit is better than a sigmoid fit to these dose-response curves and we find slopes of ΔFR = 4.6% per V/m, and ΔFR = 3% per V/m for putative pyramidal and interneurons, respectively. This difference was more pronounced at lower intensities between 100 and −100 µA (∼1.5 V/m). In this stimulation regime, putative pyramidal cells exhibited a 9.6% increase in spiking activity during cathodal TES, while putative interneurons showed a slight decrease of −0.7%. For anodal stimulation at the same intensities, putative pyramidal cells showed a decrease of −4.3%, whereas putative interneurons showed a slightly smaller decrease of −2.8% change in their firing rate. Further analysis of cell type specific effects revealed that a subset of neurons (13 out of 578 cells) responded to TES in a manner opposite to the overall average response of the population (**Fig. 3e**). Comparing the number of significantly modulated neurons across TES stimulation intensities, we found that higher intensities affected the spiking activity of more neurons regardless of the polarity of stimulation (**Fig. 3f**, mean ± SEM, n = 4 rats), and even very low intensity TES (25µA ∼ 0.375V/m) had a significant effect on the activity of a handful neurons (3.36 and 5.01% of neurons for anodal and cathodal TES, respectively).

The usual assumption is that pyramidal neurons are preferentially affected by TES due to their morphology^5^. However, the TES effects observed here appear to be rather complex, compared to what is expected from isolated stimulation of pyramidal neurons. To demonstrate this, we used transgenic mice where we can selectively stimulate excitatory cells in the CA1 region using brief pulses of blue light (405 nm, 100 ms, n = 100 trials in a head-fixed, awake transgenic mouse expressing channelrhodopsin-2 (ChR2) exclusively in CamKII expressing excitatory cells, **Suppl. Fig. 7**)^45^. We observed prominent firing of action potentials both in putative pyramidal cells and in putative interneurons, likely as a result of monosynaptic excitatory drive from the stimulated pyramidal neurons (**Suppl. Fig. 7d**, ΔFR = 95.46 and 95.24 % for putative pyramidal cells and interneurons, respectively, median, p = 0.85, Wilcoxon rank sum test).

## Discussion

First, we calibrated the recording equipment with an in vitro phantom. We also built a computational current-flow model for the rat based on high resolution MRI. This model was calibrated by measuring voltage changes in the motor cortex and hippocampus in anesthetized and freely moving rats during sinusoidal TES. We found that 100 µA currents induced 1.5-3 V/m in motor regions and 1.0-2.0 V/m in the hippocampus. Taking advantage of the 5120 contacts available on Neuropixels2.0 probes, we measured the electric fields using 1536 channels in the hippocampus. As expected from the model, electric fields decrease with distance from the stimulation electrodes. Using large-scale electrophysiology in freely moving rats, we found that neuronal firing was modulated by TES with a linear dose-response in the range of −300µA to +300 µA. Firing rate increased by about 10% per 100 µA (in **Fig. 4d**), and given an average field of approximately 1.5 V/m per 100 µA (**Fig. 3e**) this is an approximate 7% effect per V/m. This is at the upper limit of effects reported in previous in-vitro literature.^28^

Local electric field intensity and orientation at the targeted neurons is a key factor affecting the efficacy of neuromodulation^4,46^. Translation of preclinical findings is difficult because in vivo animal experiments have not measured field intensities and estimates suggest that they are ten-fold compared to humans^28^. To bridge the gap between human and animal work, and to increase rigor, it is important to know the actual magnitude and direction of field intensity in the target brain region. We recommend measuring the electric field in situ using sinusoidal waveforms at three different intensities. In order to calibrate the recording hardware^32^, we also recommend testing the stimulation and measurement systems (recording electrodes and amplifiers) in a phantom. We measured field intensity intracranially in the motor cortex and hippocampus and built a computational model to match our electrode montages. Using state-of-the-art computer models, we can now estimate the magnitude and spatial distribution of electric fields.

Currently available stimulation electrodes (saline filled cup or epicranial screw electrodes) cannot be combined with large-scale electrophysiology because of physical constraints^47^. To overcome this limitation, we developed a biocompatible permanent gel/electrode enclosure affixed to the skull and combined it with high-channel count electrophysiology and behavior in freely moving rats. This conductive gel loaded chamber provided stable current delivery to the brain and prevented chemical change at the electrode-tissue interface^48^. In many cases this is trivial to manage but with increasing invasive electrodes, higher dose, and irregular placement of electrode/electrolyte, an extreme chemical change could in theory disintegrate the skull and damage the brain.

While it is clear that the efficacy of TES depends on stimulation intensity, duration, polarity, and electrode montage (size, location, and number of electrodes)^49^, there is no reliable evidence that higher stimulation intensity is always more effective^50^. The generation of action potentials have a probabilistic nature, and the TES-induced electric fields can only bias this random process. This implies that there is no strict lower threshold for field intensity to modulate the likelihood of action potential generation. However, low intensity TES will succeed only if the neuron membrane is depolarized enough to affect firing (close to its spiking threshold). Our extracellular measurements in rats showed that even very low intensity TES (∼ 0.5 V/m) can have a significant effect on the activity of a handful of neurons (3-5 %). Previous studies have shown that affecting even a small number of neurons has significant behavioral effects^30,31,51^. We also found a dramatic increase in the number of significantly modulated neurons when the electric fields exceeded 1 V/m.

The present results may also speak to the long standing debate on the effects of endogenous electric fields on neuronal firing.^52^ Electric fields generated during theta rhythms in the hippocampus of rats^52^ can be in the range of 1-2 V/m and up to 2 V/m during slow waves in the visual cortex of ferrets^6^. New evidence that such weak fields can have an effect on neuronal function comes from in vitro experiments ^5–8^ as well as computational modeling.^53–55^ These studies mostly demonstrated a modulation of the timing of rhythmic neural activity, and relied on highly coherent rhythms that are not commonly observed in vivo. The present work extends this earlier work by demonstrating effects on firing rate for fields as low as 0.5 V/m at times scales of 2 s in vivo.

A caveat of our study is that we only analyzed acute effects on firing rate, using only short intervals of constant current stimulation (2 s). We did not aim to document lasting effects beyond the period of stimulation, although that is the primary goal of most clinical interventions with TES. A prevalent theory for long term effects of direct current TES (tDCS) is that it affects synaptic efficacy^56^. There is ample in-vitro evidence that DCS can boost synaptic plasticity^11,44,57–61^. These effects all involve an acute boost of neuronal firing in pyramidal neurons, not unlike what was observed here. Indeed, modeling studies suggest that the observed synaptic effects are due to only a small subset of active neurons^57^. Effects on synaptic plasticity have been demonstrated in-vitro down to 2.5 V/m^58^. There is no reason why the effects observed here in vivo on firing rate at 0.5 V/m would not similarly affect synaptic plasticity.

We recorded from the intermediate hippocampus because the orientation of pyramidal cells is parallel to the applied fields^23^. This ideal alignment made pyramidal cells more susceptible to electric fields. This effect was the most striking at low TES intensities. Furthermore, neurons are symmetrically located in the left and right intermediate hippocampus providing an experimental setup in which we could test cathodal and anodal effects simultaneously. Our bilateral hippocampi measurements confirmed that cathodal and anodal effects are occurring simultaneously in the two hemispheres with opposing signs (cathodal TES increased, while anodal TES decreased the spiking of neurons). When stimulation electrodes are placed on the head, it is important to consider both anodal and cathodal effects.

Neurons are embedded in networks that are influenced by TES differently. The effect of electrical stimulation is non-specific affecting any neuronal soma, and depending primarily on cell morphology relative to local field orientation.^46^ The symmetric morphology of inter-neuron suggests that their soma are not meaningfully polarized by electric fields. That they responded here similarly to pyramidal neurons is likely the result of monosynaptic drive from excitatory neurons, as we demonstrated with targeted optogenetic stimulation of pyramidal neurons. However, the spike rate increase in interneurons did not always correspond to the spike rate of monosynaptically connected pyramidal cells in the hippocampus. Indeed, pyramidal neurons on opposite hemispheres were positively affected, as expected given their cytoarchitecture. Therefore, the connectivity of individual inter-neurons may be the primary driver of how they respond to TES. A small subset of pyramidal neurons also responded opposite to other pyramidal neurons in their immediate neighborhood. As CA2 is curved it is possible that these pyramidal neurons were not aligned with the field orientation and thus their soma were minimally polarized, so that activated interneurons inhibited their firing. These findings suggest that effects on individual neurons are governed by the orientation and shape of the neuron relative to the electric field, as well as their connectivity to the network of neurons.

In conclusion, we have shown that neuronal firing rates are acutely affected in vivo at clinically relevant field magnitudes providing a viable mechanistic explanation for the effects observed with TES in human experimentation. Future work will need to establish whether these acute effects translate into long term effects, for instance, by modulating synaptic plasticity.

## Methods

### Characterization of recording and stimulation system using agar phantom

Brain phantom was constructed using a 26.7 mm diameter spherical container (30 ml syringe). To provide T1 and T2 relaxation comparable to gray matter, we followed the recipe by Schneiders at. al.^34^. A 10 mM Nickel Chloride mixture was prepared: 2.377 grams [Ni(Cl2).6H2O] per 1 L H2O * 2.377 grams of Nickel Chloride in one liter of distilled water. The agar mixture was prepared as: 3600 ml H2O, 400 ml 10 mM Ni(Cl2), 120 grams Agar, 20 grams NaCl (0.5%) and 1 gram of Sodium Azide. The mixture was heated until boiling until the agar was completely dissolved. The boiling liquid was poured into the phantom using a funnel. All air bubbles were removed by creating a vacuum in the syringe. The phantom was let cool down and a 30-mm cylinder was cut for in vitro calibration of recording and stimulation devices (**Suppl. Fig. 1a**).

High-pass filtering is inherent in the design of extracellular electrophysiology amplifiers, with bandwidths ranging from 0.1 to 10 kHz^33^. To confirm the accuracy of our recording system (RHD USB Interface Board, Intan Technologies) and determine if any signal distortion is introduced, we applied stimulation at different frequencies (1, 10, 100 and 1000 Hz) and at different intensities (100, 150 and 200 µA, **Suppl. Fig. 1a**) to an agar phantom^34,35^. The phantom was a homogeneous cylinder of 20 mm in height and 26.7 mm in diameter that was filled with agar with conductivity σ = 0.9 S/m (**Suppl. Fig. 1b**). Stimulation was delivered using platinum electrodes (2.2 by 1.6 mm) positioned at a separation of 17.81 mm using an isolated stimulus generator (STG 4002, Multichannel Systems). For the measurement of the voltage values generated during stimulation within the phantom, we used two custom-built tungsten electrodes (two recording channels each electrode, 56.3 ± 19.8 kOhm impedance at 1 kHz, mean ± SD, **Suppl. Fig. 1b, c and Suppl. Fig. 2 and Suppl. Video 1**). The tungsten electrodes were attached to a microdrive^36^ and positioned 3.4 mm apart using a stereotactic frame (Model 962, David Kopf Instruments, **Suppl. Fig. 1b and d**). The magnitude of the electric field increased linearly with stimulation intensities as expected (100, 150 and 200 µA, **Suppl. Fig. 1e**). However, the slope of the electric field decreased during 1 Hz stimulation (**Suppl. Fig. 1e**) reflecting signal attenuation caused by the built-in 0.7 Hz high-pass filter in the recording system.

### Preparing tungsten recording device

A 26-gauge needle was cut to 3 mm. 50-µm tungsten wires (Tungsten 99.95%, 100211, insulated with Heavy Polyimide, HML – Green, California Fine Wire, CA) were cut to 30 mm and the insulation (green coating) was removed from one end using a razor blade. Two tungsten wires were inserted into the stainless-steel tube (2-channel shank). Wires were positioned 5 mm from the end of the tube. Wires were separated (ch-1 and ch-2) 1 mm apart from each other (**Suppl. Fig. 3**). Ultra-liquid superglue (Loctite 1647358, Henkel, Germany) was applied on both ends of the tube and between wires. Two, 2-channel, single shank devices were attached to a mechanical shuttle (microdrive^36^) or a 2 by 4 mm printed circuit board) making a 4-channel, 2-shank device. For the motor cortex recording wires were bent 90 degrees. Tungsten wires and a ground wire were soldered inside a header pin (575-8514305010, Mouser, TX). The header pin connector to Omnetics adapter was soldered to connect tungsten wires to preamplifier headstage (#C3324, Intan Technologies Inc., CA). Impedance of the wires were measured by RHD USB interface board from Intan (Intan Technologies LLC, CA, USA). The device was lowered into 0.9% saline and connected to the recording preamplifier ground (RHD 32-channel recording headstages). Impedance measurement was performed at 1 kHz frequency.

### Experiments on rats

All experiments were approved by the Institutional Animal Care and Use Committee at New York University Medical Center and CUNY IACUC. Rats (adult male n = 6 and female n = 1, 300–400 g) were kept in a vivarium on a 12-hour light/dark cycle and were housed two per cage before surgery. Rats were implanted with custom-made recording and stimulating electrodes under urethane anesthesia (1.3–1.5 g/kg, intraperitoneal injection). Atropine (0.05 mg kg–1, s.c.) was administered after anesthesia induction to reduce saliva production. The body temperature was monitored and kept constant at 36–37 °C with a DC temperature controller (TCAT-LV; Physitemp, Clifton, NJ, USA). Stages of anesthesia were maintained by confirming the lack of a nociceptive reflex.

### Recording electric fields in motor cortex of anesthetized rats

The chest wall and the head were shaved. We made an incision on the head and on the chest wall. A 10 by 10 mm platinum mesh electrode (Goodfellow, PT00-MS-000110) was sutured to the pectoral muscle and an insulated cable was tunneled to the top of the head of the animal^19^. The skull was cleaned by hydrogen peroxide (2%) and a stimulation pocket was attached to the skull using dental cement (1.5 mm anterior to bregma and 3 mm lateral to midline). The pocket was filled with conductive gel (Signagel Electrode Gel) and a 3 by 3 mm platinum stimulation electrode was inserted inside. A craniotomy was performed on the temporal bone (1.44 mm anterior from bregma and 3 mm deep from the top of the skull) and the dura was removed. The tungsten device was inserted to the target depth (2.4 mm from the surface of the brain). The collected data was digitized at 20 kS/s using an RHD2000 recording system (Intan Technologies, Los Angeles, CA). Stimulation was delivered by Caputron LCI 1107 High Precision. Varying frequencies (10, 100 and 1000 Hz) at varying intensities (10, 20 and 40 µA) were delivered through the stimulating electrodes. Electric field was measured by fitting a sinusoid to the recorded voltage differences between the 4 contacts, averaging amplitudes of the two parallel measures, and dividing by the electrode distance (1 mm). This results in a 2D field vector, with magnitude given by the norm of this vector.

### Recording electric fields in hippocampus of anesthetized and freely moving rats

The skin of the head was shaved. After a midline incision the surface of the skull was cleaned by hydrogen peroxide (2%). A custom stimulation pocket was attached to the skull using dental cement (4.8 mm posterior from bregma). The pocket was filled with conductive gel (SuperVisc, EasyCap GmBH, Germany) and a 2 by 2 mm platinum stimulation electrode (#349356-600MG, Sigma-Aldrich, Inc., St. Louis, MO) was inserted inside. A stainless-steel ground screw was placed above the cerebellum (#90910A380, McMaster-Carr, Elmhurst, IL). A craniotomy was performed (4.8 mm posterior from Bregma and 5 mm lateral to midline) and the dura was removed. The silicon probe was attached to a microdrive^36^ (128-5, Diagnostic Biochips Inc., Glen Burnie, MD or Neuropixels 2.0) and it was inserted to the target depth (4 and 6 mm from the surface of the brain). We constantly monitored the electrophysiological signal during insertion. The collected data (128-5 probe) was digitized at 20 kS/s using an RHD2000 recording system (Intan Technologies, Los Angeles, CA). Neuropixels2.0 data was digitized at 30 kS/s and a custom PXIe (Peripheral Component Interconnect (PCI) eXtension for Instrumentation; a standardized modular electronic instrumentation platform) data acquisition card was connected to a computer via a PXI chassis (NI 1071, National Instruments, Austin, TX), and OpenEphys software was used to write the data to disk^43,62^. Baseline session (one hour before TES) and electrical stimulation session were recorded in the homecage of rats during the sleep cycle of the animals. Stimulation was delivered by an STG4002–16mA (Multi Channel Systems, Reutlingen) using different intensities and polarities (**Suppl. Table 1**). Rats did not show any behavioral response to stimulation. To measure the electric fields in the hippocampus, varying frequencies (10, 100 and 1000 Hz) at varying intensities (10, 20 and 40 µA) were delivered through the stimulating electrodes at the end of the recording session. TES induced voltage changes were measured shank-by-shank (4*384 = 1536 recording sites in total). Electric field was measured by fitting a sinusoid to the recorded voltage at each recording site. We first calculated the average peak-to-peak voltage on each site (n = 500 trials), and then calculated the first spatial derivative of these voltage values across shanks. An average hippocampal electric field was calculated after localizing the cellular layer of the hippocampus using electrophysiological markers (Fig. 3, n = ± 32 channels were averaged around the center of the pyramidal layer).

### Local Field Potential Analysis

To detect sharp wave ripples a single electrode in the middle of the pyramidal layer was selected. The wide-band LFP signal was band-pass filtered (difference-of-Gaussians; zero-lag, linear phase FIR), and instantaneous power was computed by clipping at 5 SD, rectified and low-pass filtered. The low-pass filter cut-off was at 55 Hz, and the band-pass filter was from 80 to 200 Hz. Subsequently, the power of the non-clipped signal was computed, and all events exceeding 5 SD from the mean were detected. Events were then expanded until the (non-clipped) power fell below 2 SD; short events (<15 ms) were discarded. The pyramidal layer of the CA1 region was identified physiologically by increased unit activity and characteristic LFP patterns.

### Single unit analysis

A concatenated signal file was prepared by merging all recordings from a single animal from a single day. To improve the efficacy of spike sorting, stimulation induced onset and offset artefacts were removed before automatic spike sorting (10 ms before and 100 ms after the detected artefacts, linear interpolation between timestamps). Putative single units were first sorted using Kilosort^63^ and then manually curated using Phy (https://phy-contrib.readthedocs.io/). After extracting timestamps of each putative single unit activity, peristimulus time histograms and firing rate gains were analyzed using a custom MATLAB (Mathworks, Natick, MA) script. Changes in firing rate of single units (ΔF) were calculated by the following equation:

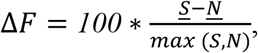

Where *SS* and *NN*, are the mean firing rates for the stimulation (S) and no stimulation (N) epochs. Cells were classified into three putative cell types: narrow interneurons, wide interneurons, and pyramidal cells based on waveform metric^42^.

### Cell Type Classification

In the processing pipeline, cells were classified into two putative cell types: interneurons, and pyramidal cells. Interneurons were selected by two separate criteria. We labeled single units as interneurons if their waveform trough-to-peak latency was <0.425 ms, or if the waveform trough-to-peak latency was >0.425 ms and the rise time of the autocorrelation histogram was >6 ms. The remaining cells were assigned as pyramidal cells. Autocorrelation histograms were fitted with a triple exponential equation to supplement the classical, waveform feature based single unit classification (https://cellexplorer.org/pipeline/cell-type-classification/) ^42^. Bursts were defined as groups of spikes with interspike intervals < 9 ms. The authors had isolated 762 putative single units from seven animals in nine sessions (n = 453 putative pyramidal cells, n = 193 putative interneurons).

### Detection of Monosynaptic Cell Pairs

Cross-correlation (CCG) analysis has been applied to detect putative monosynaptic connections^64,65^. CCG was calculated as the time resolved distribution of spike transmission probability between a reference spike train and a temporally shifting target spike train. A window interval of [−5, +5] ms with a 1-ms bin size was used for detecting sharp peaks or troughs, as identifiers of putative monosynaptic connections. Significantly correlated cell pairs were identified using a previously ground-truth validated convolution method^64^. The reference cell of a pair was considered to have an excitatory monosynaptic connection with the referred neuron, if any of its CCG bins within a window of 0.5–3 ms reached above confidence intervals.

### Modeling of current-induced fields

Magnetic resonance imaging (MRI) scan of a template rat head was segmented into nine tissue masks namely scalp, skull, cerebrospinal fluid (csf), gray matter, white matter, cerebellum, hippocampus, thalamus, and air to develop a high resolution (0.1 mm) MRI derived volume conductor model in Simpleware (Synopsys Inc., CA, USA) using both automatic and manual filters. Computer aided model (CAD) geometry of the electrodes were modeled in SolidWorks (Dassault Systemes Corp., MA, USA) and positioned based on coordinates value from the experiment. Specifically, we modeled two montages to predict the electric field in the motor cortex (montage 1) and hippocampus (montage 2). In montage 1, Platinum electrode (anode: 3 x 3 x 0.1 mm3) was positioned above the primary motor cortex over the exposed skull by smearing a thin layer of conductive electrode gel, whereas the return electrode (Platinum mesh) was placed inside the chest wall (cathode: 10 x 10 x 1 mm3). In montage 2, a Platinum electrode (anode: 2 x 2 x 0.1 mm3) was immersed into a conductive electrode gel and secured over the temporal bone by a plastic electrode holder on each hemisphere of the rodent head.

An adaptive tetrahedral mesh of rat model resulting from multiple mesh refinements was generated using a voxel-based meshing algorithm and contained > 8 M tetrahedral elements and was solved for > 10 million degrees of freedom. Volumetric meshes were later imported into COMSOL Multiphysics 4.3 (COMSOL Inc., MA, USA) to solve the model computationally using a steady-state assumption (Laplace equation, ∇(σ∇V) =0, where V= potential and σ = conductivity^37^). Compartment-specific assigned electrical conductivities were given as, scalp: 0.465 S/m; skull: 0.01 S/m; csf: 1.65 S/m; air: 1×10-15; gray matter: 0.276 S/m; cerebellum: 0.276 S/m; hippocampus: 0.126 S/m; white matter: 0.126 S/m; thalamus: 0.276 S/m, electrode: 5.99 x 107 S/m, conductive gel: 4.5 S/m, and plastic electrode holder 1×10-15 S/m. All values were based on prior literature^66,66^. The boundary conditions were applied as current (Montage 1: 150 µA and Montage 2: 80 µA) at the exposed surface of the anode while the contralateral electrode was grounded (cathode). All remaining outer boundaries of both models were electrically insulated. Electric field at the primary motor cortex and hippocampus, mimicking experimental recording sites, was predicted and peak value was reported.

### Statistical Analysis

Statistical analyses were performed with MATLAB functions or custom-made scripts. The unit of analysis was typically identified as single neurons. In a few cases, the unit of analysis was sessions or animals, and this is stated in the text. Unless otherwise noted, non-parametric two-tailed Wilcoxon rank-sum (equivalent to Mann-Whitney U-test) or Wilcoxon signed-rank test was used. On box plots, the central mark indicates the median, bottom and top edges of the box indicate the 25th and 75th percentiles, respectively, and whiskers extend to the most extreme data points not considered outliers. Outliers are not displayed in some plots but were included in statistical analysis. Due to experimental design constraints, the experimenter was not blind to the manipulation performed during the experiment (transcranial electrical stimulation manipulation).

## Data availability

The data sets generated and analyzed during the current study are available upon reasonable request from the corresponding authors for further analyses.

## Acknowledgements

We thank Gyorgy Buzsaki for useful comments on the manuscript. This work was supported by NIH through grant R01 NS130484.

## Author contributions

FF, MV performed surgeries, collected, and analyzed data, NK performed current flow simulations. FF, NK, LCP, MB and MV wrote the manuscript.

## Competing interests

LP is listed as inventor in patents owned by CCNY, and has shares in Soterix Medical Inc. The City University of New York (CUNY) has IP on neuro-stimulation systems and methods with authors NK and MB as inventors. NK is an employee of Synchron Inc and consults for Ceragem Medical. MB has equity in Soterix Medical. MB consults, received grants, assigned inventions, and/or served on the S A B of SafeToddles, Boston Scientific, GlaxoSmithKline, Biovisics, Mecta, Lumenis, Halo Neuroscience, Google-X, i-Lumen, Humm, Allergan (Abbvie), Apple, Ybrain, Ceragem Medical, Remz.

