## Supplemental File for "Transcranial electric stimulation modulates firing rate at clinically relevant intensities"

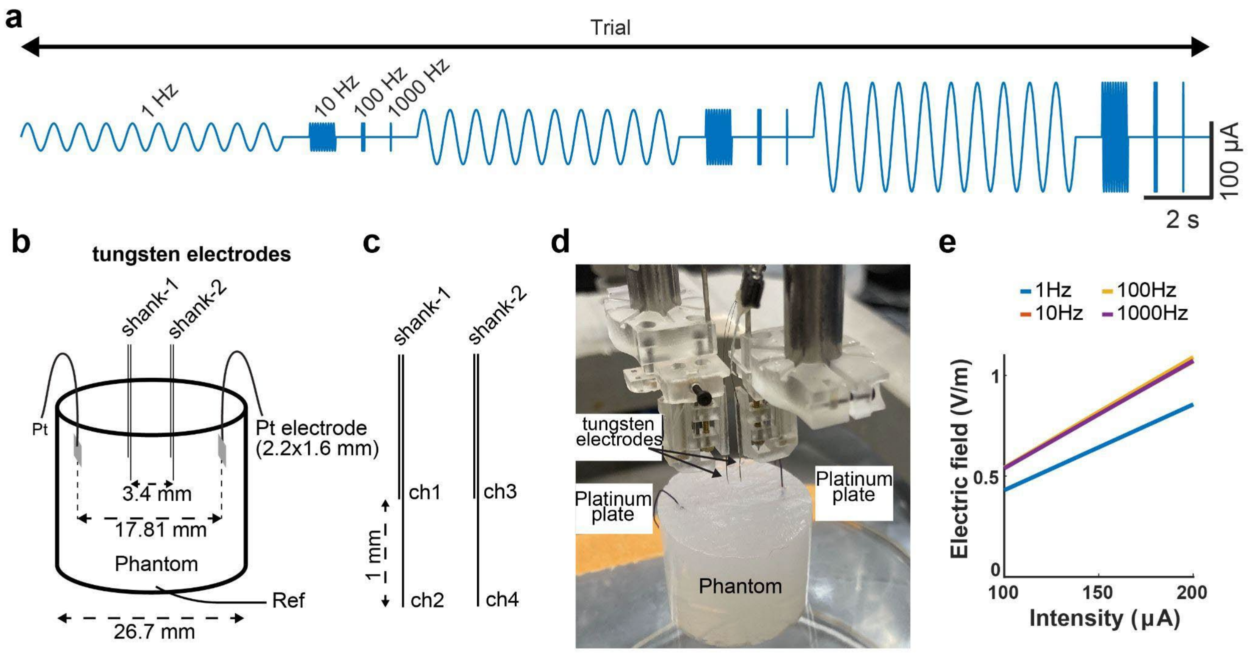


**Supplementary Figure 1. Validating the recording and stimulation system in a brain phantom. a)** Schematic of stimulation waveform. 4 frequencies (1, 10, 100 and 1000 Hz) at 3 different intensities (100, 150 and 200 µA) were used. A single trial consisted of 10 cycles of each condition separated by 1 second of stimulus free interval (n = 30 trials, 300 cycles/frequency/intensity). **b)** Schematic of recording setup. Platinum stimulation electrodes (2.2 by 1.6 mm) were inserted into the phantom (3 mm from the surface of the phantom). The stimulation electrodes were separated by 17.81 mm. Two, 2-channel recording electrodes (50 µm tungsten wires) were mounted on microdrives and were inserted into the phantom using a stereotactic frame (2.65 mm and 3.17 mm from the surface). The distance between the recording electrodes was 3.4 mm. A stainless-steel reference wire was inserted at the bottom of the phantom. The stimulation signal was generated with an STG-4002 isolated signal generator. The voltage signals were recorded with an Intan USB Eval Board and were digitized at 20 kS/s. **c)** Schematic of the recording devices. The contact sites were separated by 1 mm. **d)** Photograph of the recording setup. Note that the phantom was placed on a glass baker to isolate it from the stereotactic frame. **e)** The electric field increases linearly with increasing stimulation intensity (10 Hz slope = 5.31 V/m/µA; 100 Hz slope = 5.48 V/m/µA, 1000 Hz slope = 5.34 V/m/µA). Note the difference of 1 Hz compared to the rest of the frequencies (1 Hz slope = 4.26 V/m/µA).

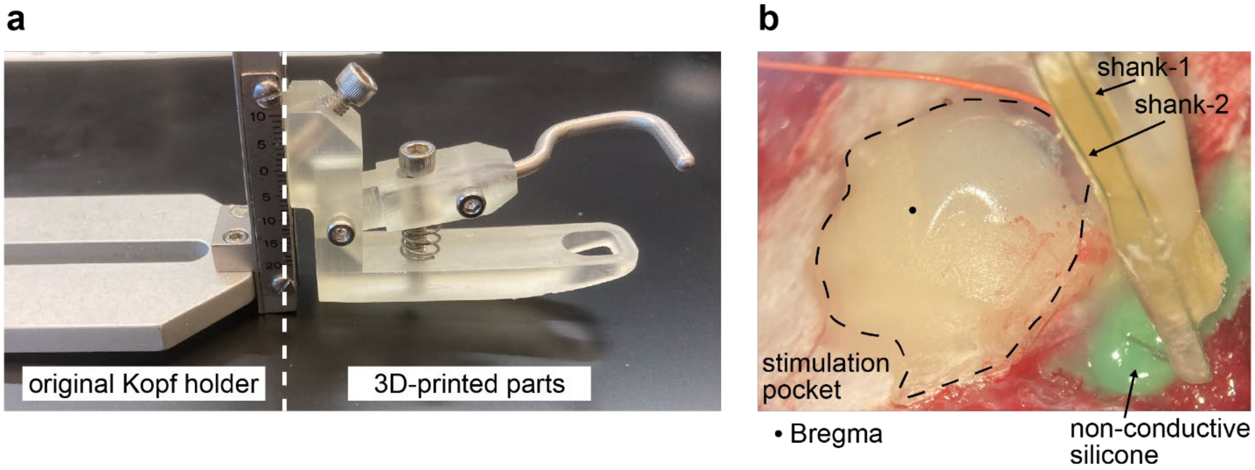


**Supplementary Figure 2. Measuring intracerebral electric field magnitude in the motor cortex of anesthetized rats. a)** A 3D-printed nose holder is combined with Kopf stereotactic holder. **b)** Intraoperative photograph of the measurement of intracerebral electric fields by a 2-shank, 4-channel electrode in an anesthetized rat. The recording electrode is inserted through the temporal bone and the craniotomy is sealed with non-conductive silicone (green silicone). Stimulation pocket (outlined by black dashed line) is attached to the skull using dental cement (black dot shows the location of Bregma). The pocket is filled with conductive gel and the stimulation electrode is placed inside the pocket (platinum plate is attached to the orange stimulation cable). The pocket is sealed with dental cement.

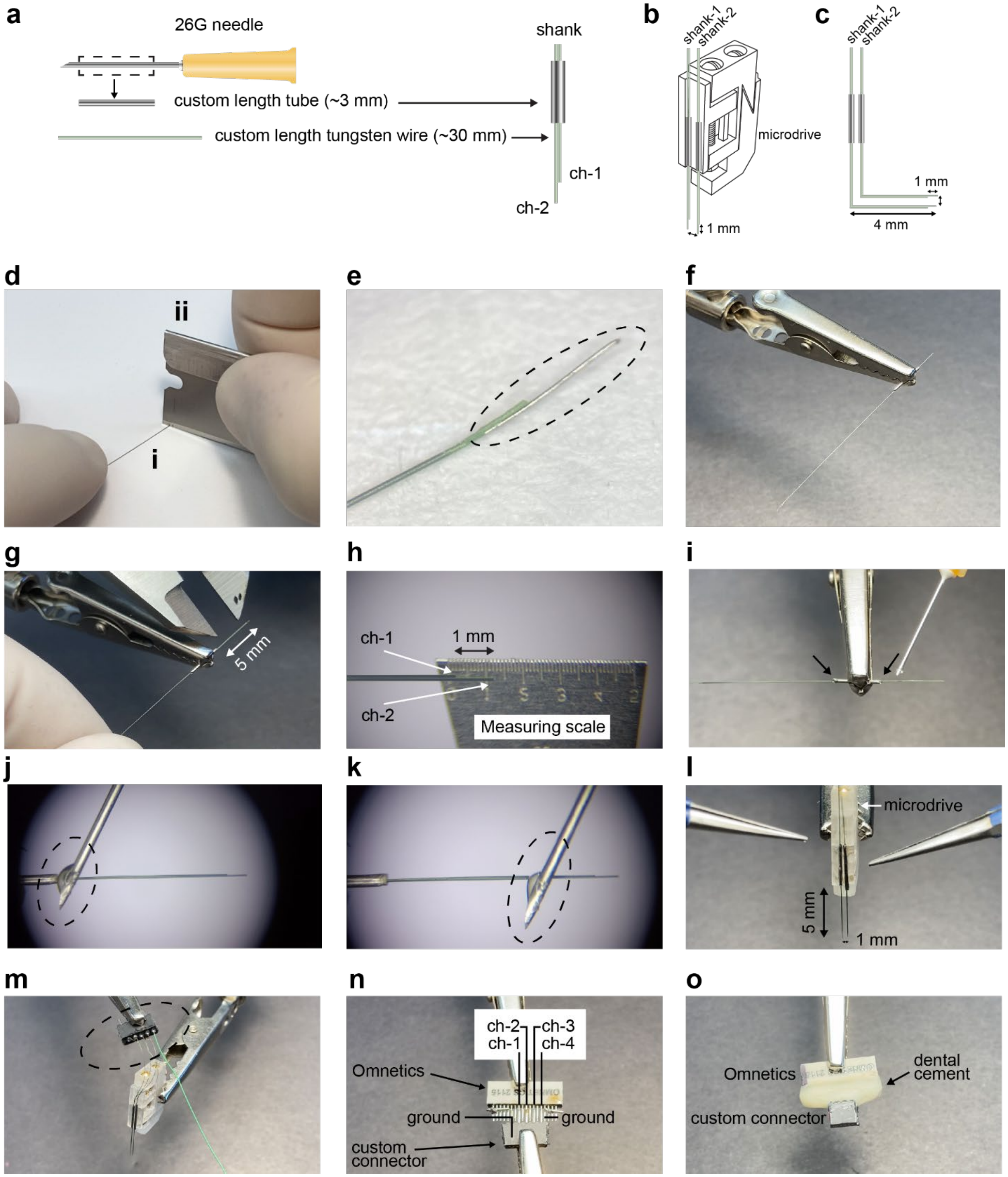


**Supplementary Figure 3. Preparation of tungsten recording electrode. a)** A 26-gauge needle serves as a stainless-steel tube, and it is cut to a custom length (3 mm). 50-µm tungsten wires are also cut to a custom length (30 mm). Right: two tungsten wires are inserted into the stainless-steel tube making a 2-channel (ch-1 and ch-2) on a single shank device. **b)** Two, 2-channel, single shank devices are attached to a mechanical shuttle (microdrive) making a 4-channel, 2-shank device. **c)**

Custom modification (90 degrees bend) for motor cortex recording. Tungsten wires can be bent. **d)** Removal of insulation (green coating) from one end of the tungsten wire (**i**) using a razor blade (**ii**). **e)** Dashed line shows the uninsulated segment of the tungsten wire. **f)** Place wires inside a custom-cut 26G needle (~4 mm tube). **g)** Position the wires to the desired distance using a caliper (5 mm from the end of the tube). **h)** Separate wires (ch-1 and ch-2) to desired distance using a measuring scale (1 mm apart from each other). **i)** Apply ultra-liquid superglue on both ends of the tube (black arrows). **j)** Application of superglue at high magnification (dashed line shows a blob of superglue at the tip of a 26G needle). **k)** Apply ultra-liquid superglue between wires. Make sure to arrange wires in parallel, without any distance between them. **l)** Build the final configuration (1 mm distance between shanks) and attach the tubes to a holder (microdrive). **m)** Solder tungsten wires and a ground wire inside the connector (dashed line). **n)** Align header pin connector with Omnetics and solder them together. Unused pins should be shorted to ground. **o)** Cover soldering with dental cement.

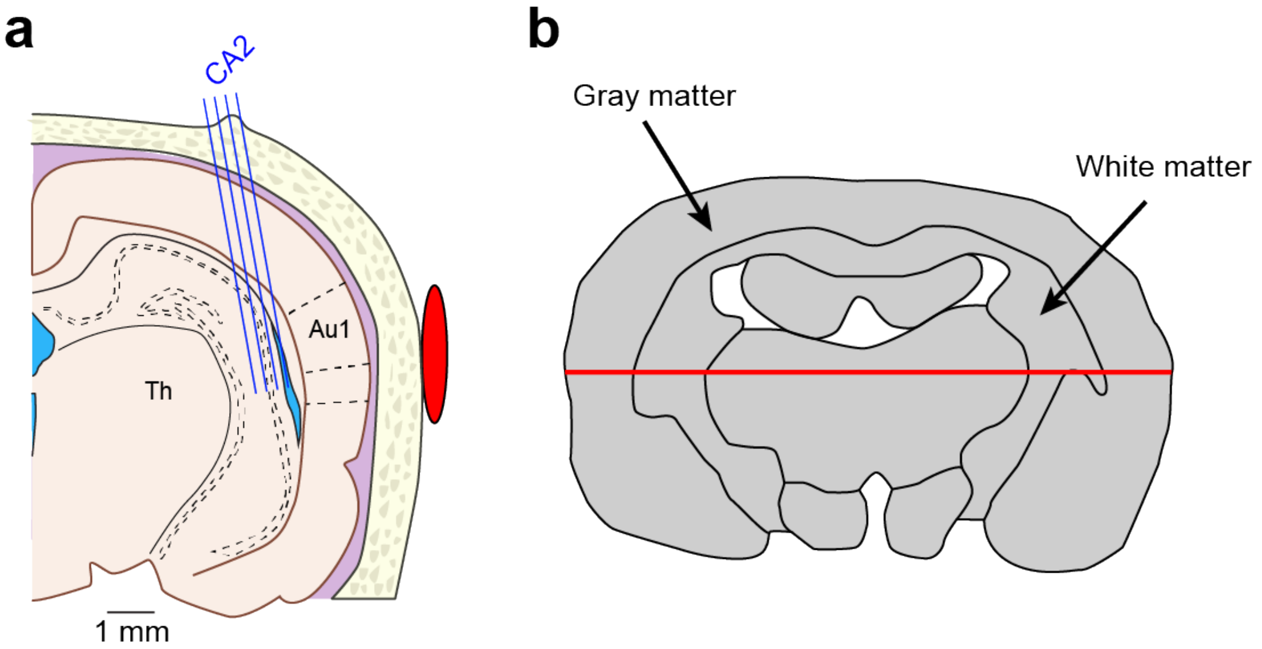


**Supplementary Figure 4. Gray and white matter have different conductivity. a)** Schematic of recording shank location in the hippocampus. **b)** The difference in conductivity between gray and white matter results in an electric field discontinuity. The red line corresponds to the white dotted horizontal line as in **Fig. 3g.**

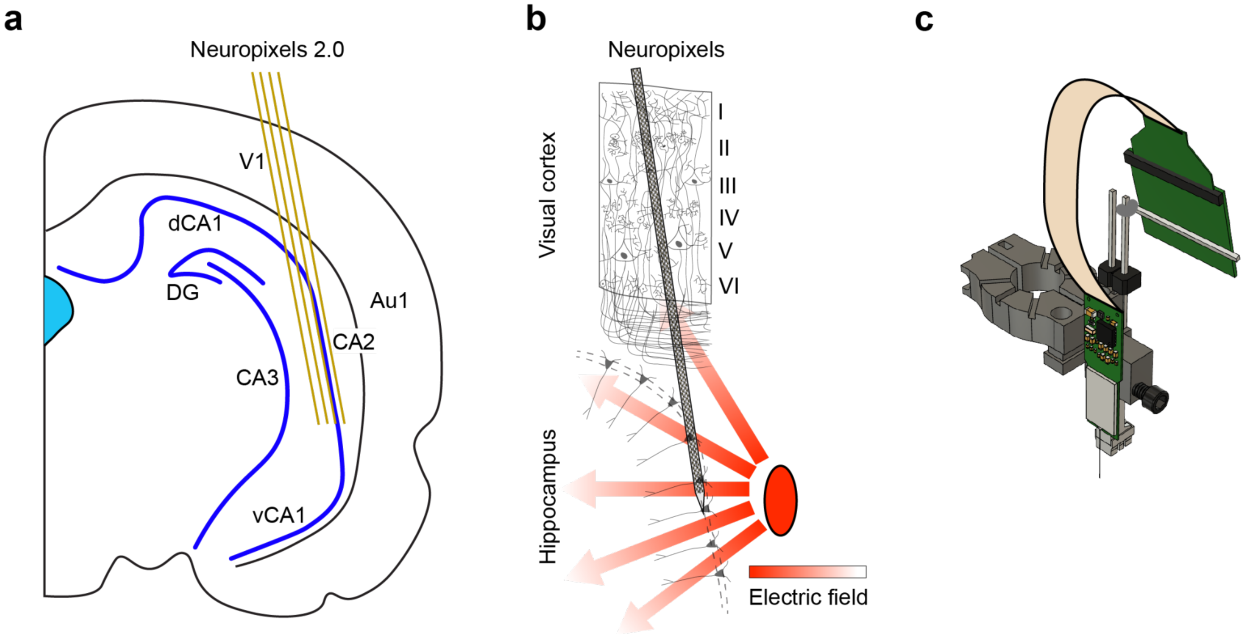


**Supplementary Figure 5. Experimental setup in freely moving rats a)** Schematic of experimental setup. Hippocampal single units were recorded from the intermediate CA2 (4.8 mm posterior to bregma and 5.6 mm lateral to midline, 10 degree angle). V1-primary visual cortex, Au1-primary auditory cortex, dCA1-dorsal CA1, vCA1-ventral CA1, DG-dentate gyrus. **b)** Schematic of orientation of neurons relative to the electric field in hippocampus (CA2) and visual cortex. CA2 pyramidal cells are aligned parallel to the electric fields. **c)** Neuropixels probe is attached to a metal microdrive (probe, microdrive and implantation tool are shown).

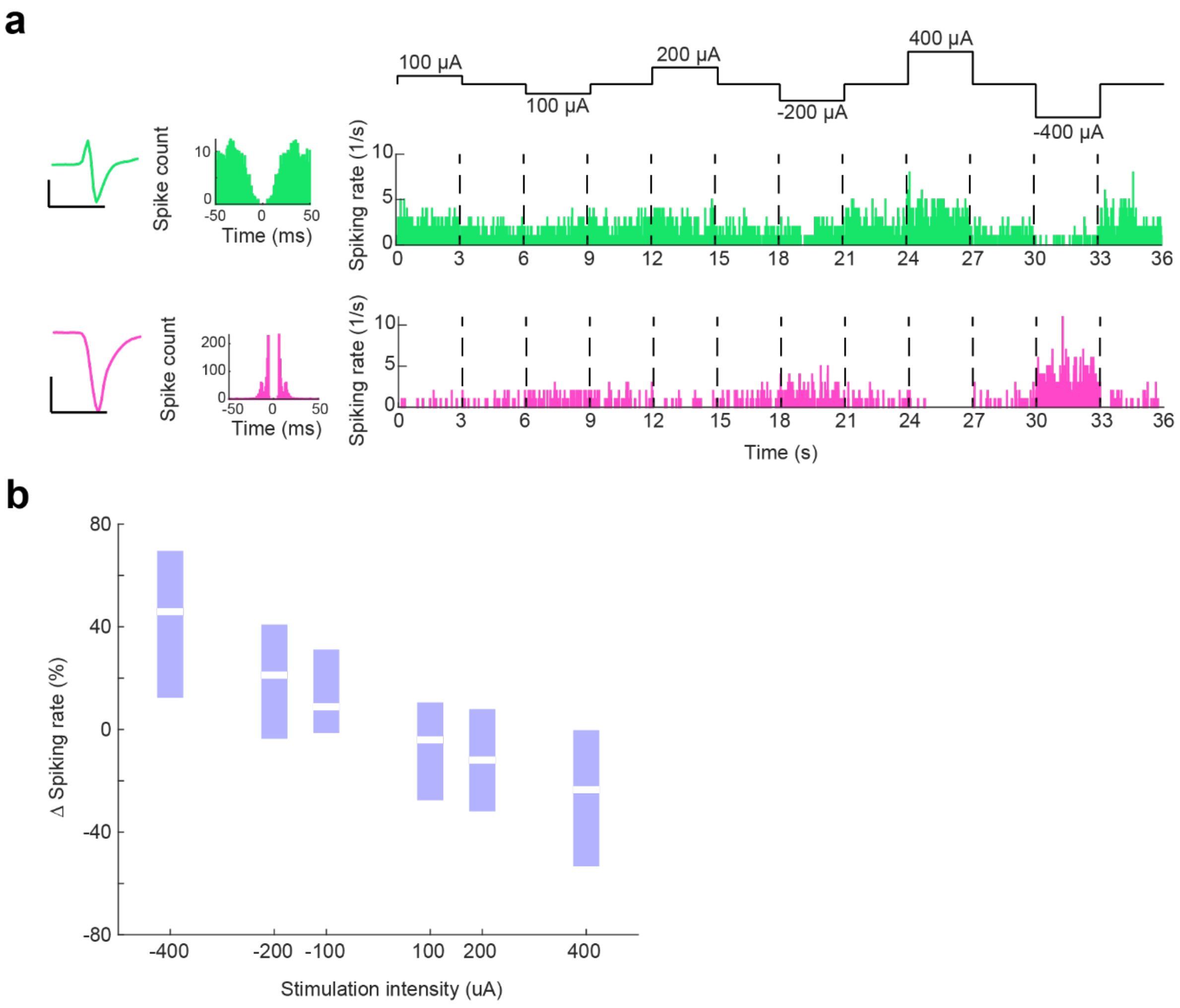


**Supplementary Figure 6. Single unit effects of high intensity TES in a urethane anesthetized rat. a)** Response of two example neurons. The putative interneuron (top, mean waveform and autocorrelation histogram is shown on the left) was strongly inhibited while the putative pyramidal cell (bottom, mean waveform and autocorrelation histogram is show on the left) was strongly excited by cathodal TES (intensity was -400 µA, 3s stimulation was followed by 3s stimulation free epochs, n = 89 trials), as shown by peristimulus time histograms (right panels). **b)** Changes of spiking activity in response to TES (±100, ±200 and ±400 µA, 3s On, 3s Off, n = 89 trials). Note linear changes of spiking rate with changing polarity and amplitude of external electric fields (slope: 4.15% per V/m, R = -0.52, p < 0.001, n = 68 neurons).

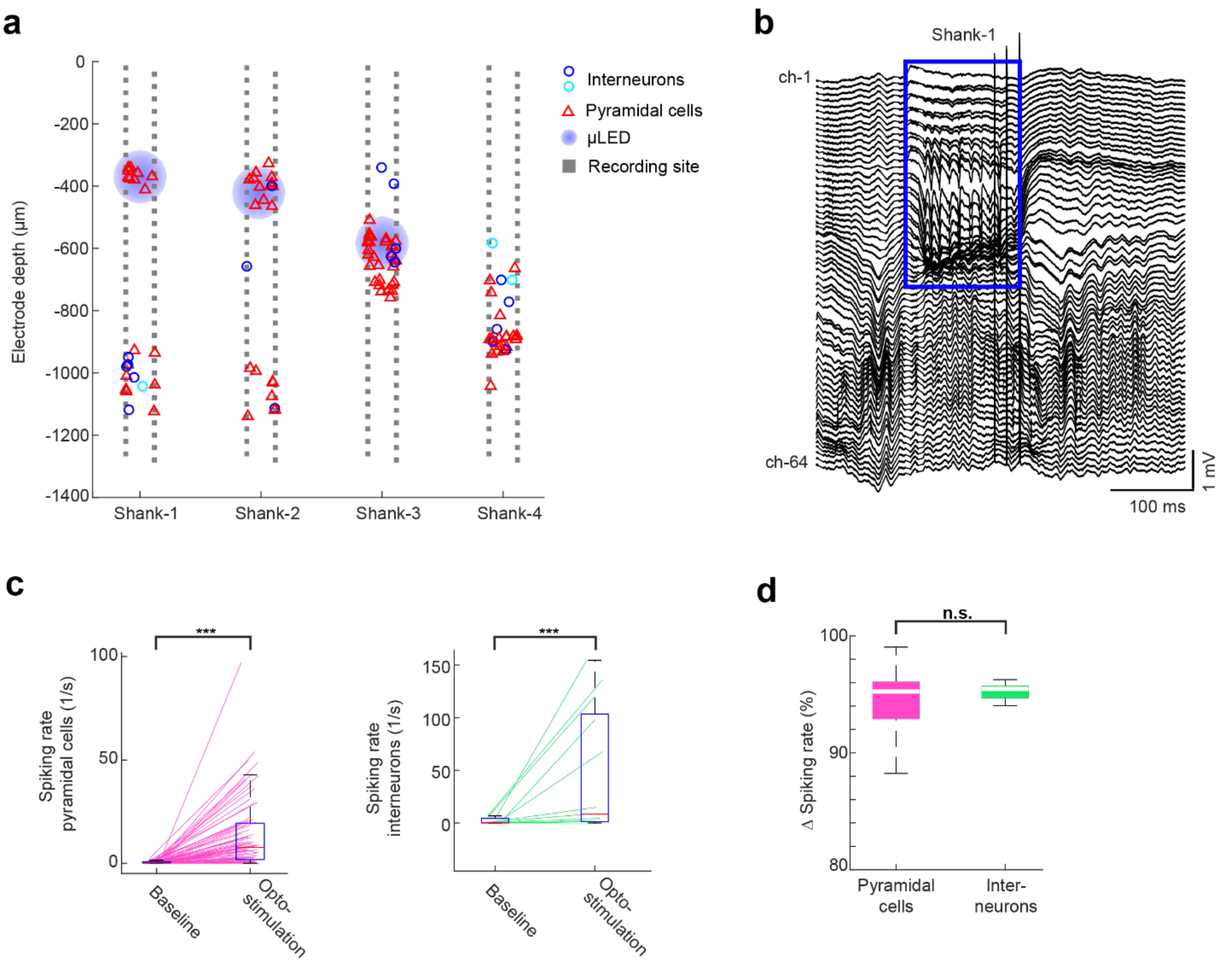


**Supplementary Figure 7. Single unit effects of µLED optogenetic stimulation in a head-fixed, awake mouse. a)** Probe layout is shown with the putative location of recorded neuron somata of putative pyramidal cells and interneurons (red triangle and blue, cyan circle, respectively). Single units were clustered in the cellular layers of hippocampus (0 µm represents dorsal channels on the probe). Optogenetic stimulation was delivered using a blue µLED located on the shanks (shank-1 to 3, blue circles). **b)** Depth profiles of LFPs show an example response elicited by a 150 ms µLED activation in the CA1 region (blue globes in **a**). Note the stimulation effect is restricted to the CA1 region only (blue rectangle). **c)** Left: the spiking rate of putative pyramidal cells increased from 0.68 ± 0.098 Hz to 15.41 ± 2.17 Hz (n = 65 neurons, mean ± SEM, p < 0.01, Wilcoxon rank sum test). Right: monosynaptic connections between pyramidal cells and interneurons (24 putative pyramidal cells connected to 8 putative interneurons) led to an increase in the spiking rate of putative interneurons from 2.53 ± 0.84 Hz to 55.33 ± 18.48 Hz (n = 11 neurons, mean ± SEM, p < 0.01, Wilcoxon rank sum test). **d)** Gain in spiking activity was similar between excitatory cells and inhibitory neurons (ΔFR = 95.46 % in putative pyramidal cells vs. 95.24 % in putative interneurons, median, p = 0.85, Wilcoxon rank sum test).

| Session name | Sex | Probe | Stimulus intensity (μA) | Stimulus durations (s) |
|---|---|---|---|---|
| Rat_01 | F | NP1 | ±100, ±200, ±300 | 4 |
| Rat_02 | M | 128-8, dual sided, DBC | ±25, ±50, ±100, ±200, ±300 | 3 |
| Rat_03 | M | NP2.0 | ±25, ±50, ±100, ±200, ±300 | 3 |
| Rat_04 | M | NP2.0 | ±25, ±50, ±100, ±200, ±300 | 3 |
| Rat_05* | M | Buz32, NeuroNexus | 200 | 0.5 |
| Rat_06* | M | 128-8, dual sided, DBC | ±100, ±200, ±400 | 4 |

**Supplementary Table 1.** Stimulation parameters used during electrophysiology experiments. Star indicates sessions under urethane anesthesia.

**Supplementary Video 1. Preparation of tungsten recording electrode.**
